# Controlled cycling and quiescence enables homology directed repair in engraftment-enriched adult hematopoietic stem and progenitor cells

**DOI:** 10.1101/301176

**Authors:** Jiyung Shin, Stacia K. Wyman, Mark A. Dewitt, Nicolas L Bray, Jonathan Vu, Jacob E. Corn

## Abstract

Hematopoietic stem cells (HSCs) are the source of all blood components, and genetic defects in these cells are causative of disorders ranging from severe combined immunodeficiency to sickle cell disease. However, genome editing of long-term repopulating HSCs to correct mutated alleles has been challenging. HSCs have the ability to either be quiescent or cycle, with the former linked to stemness and the latter involved in differentiation. Here we investigate the link between cell cycle status and genome editing outcomes at the causative codon for sickle cell disease in adult human CD34+ hematopoietic stem and progenitor cells (HSPCs). We show that quiescent HSPCs that are immunophenotypically enriched for engrafting stem cells predominantly repair Cas9-induced double strand breaks (DSBs) through an error-prone non-homologous end-joining (NHEJ) pathway and exhibit almost no homology directed repair (HDR). By contrast, non-quiescent cycling stem-enriched cells repair Cas9 DSBs through both error-prone NHEJ and fidelitous HDR. Pre-treating bulk CD34+ HSPCs with a combination of mTOR and GSK-3 inhibitors to induce quiescence results in complete loss of HDR in all cell subtypes. We used these compounds, which were initially developed to maintain HSCs in culture, to create a new strategy for editing adult human HSCs. CD34+ HSPCs are edited, allowed to briefly cycle to accumulate HDR alleles, and then placed back in quiescence to maintain stemness, resulting in 6-fold increase in HDR/NHEJ ratio in quiescent, stem-enriched cells. Our results reveal the fundamental tension between quiescence and editing in human HSPCs and suggests strategies to manipulate HSCs during therapeutic genome editing.

## Introduction

Hematopoietic stem cells (HSCs) ensure the lifelong production of all blood cells through their unique capacity to self-renew and to differentiate (Figure 1A). Deficiencies in HSC renewal lead to severe anemias such as Fanconi Anemia and Diamond Blackfan Anemia (Corey et al., 2007). Inappropriate differentiation can lead to either the over- or under-production of blood components, causing disorders that range from immunodeficiency to cancer.

**Figure 1.**
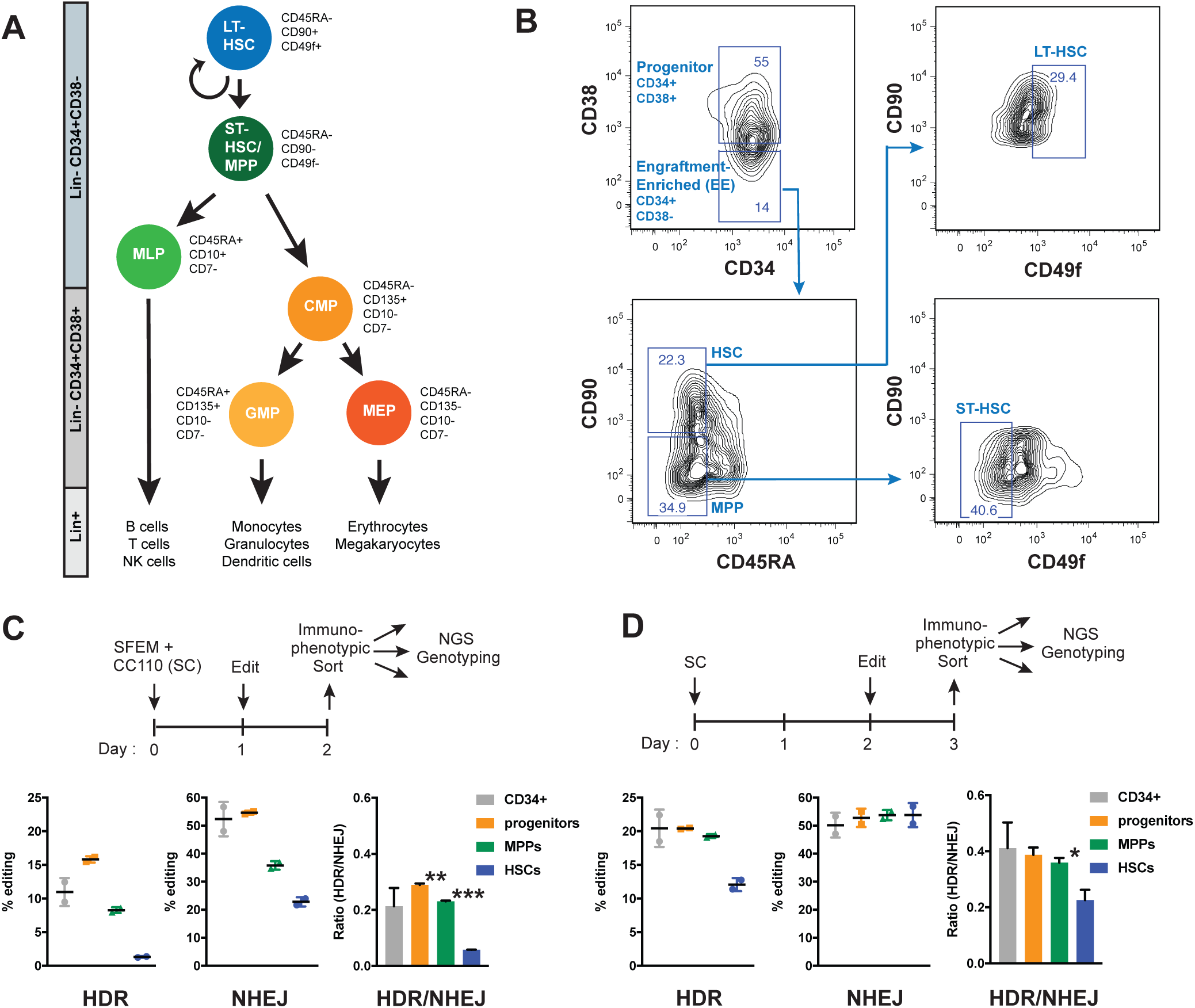
HSCs require more time to activate HDR pathways during gene editing as compared to differentiated cells. (A) Diagram describing the human hematopoietic population hierarchy. long-term hematopoietic stem cell (LT-HSC); short-term hematopoietic stem cell (ST-HSC); multipotent progenitor (MPP); multi-lymphoid progenitor (MLP); common myeloid progenitor (CMP); granulocyte monocyte progenitor (GMP); megakaryocytic-erythroid progenitor (MEP). Immunophenotypic markers for each subpopulation adapted from (Notta et al., 2011). (B) Fluorescence-activated cell sorting scheme to isolate human progenitors (CD34+ CD38+), Engraftment-enriched (EE) HSPCs (CD34+ CD38-), MPPs, HSCs, ST-HSCs, and LT-HSCs from human mobilized peripheral blood CD34+ HSPCs. CD34+ cells were stained with monoclonal antibodies against CD34, CD38, CD45RA, CD90 (Thy1), and CD49f antigens. The frequency of each subpopulation is based on the parent gate. (C) Editing outcomes in CD34+ subpopulations 1 day post electroporation, 2 days in culture. Percentage of reads positive for HDR or NHEJ by next-generation amplicon sequencing at the HBB site. HSCs lack HDR alleles. Representative data from n≥3 independent experiments with n≥2 biological replicates for each condition. Mean ± SD shown. **: p<0.01, ***: p<0.001 by unpaired t-test. (D) Editing outcomes in CD34+ compartments 1 day post electroporation, 3 days in culture. Percentage of reads positive for HDR or NHEJ by next-generation amplicon sequencing at the HBB site. HSCs accumulate significant HDR alleles although they show lower HDR/NHEJ ratio compared to MPPs and progenitors. Representative data from n≥3 independent experiments with n≥2 biological replicates for each condition. Mean ± SD shown. **: p<0.05 by unpaired t-test. *See also Figure S1*.

Due to their ability to simultaneously self-renew and generate the entire blood system, long-term HSCs represent an attractive target for gene therapy and editing to deliver lasting treatments for blood disorders. Genome editing, most recently accelerated by the development of CRISPR-Cas reagents, has emerged as an effective tool to precisely target human HSCs (Dever et al., 2016; DeWitt et al., 2016; Genovese et al., 2014b; Hoban et al., 2015; De Ravin et al., 2017; Wang et al., 2015). However, the replacement of genetic sequences via nuclease-induced HDR in HSCs has lagged behind the ability to disrupt sequences via NHEJ. Specifically, while bulk-edited CD34+ populations of HSPCs exhibit high levels of HDR after a few days in culture, sustained engraftment of HDR-edited cells in the Long-Term engrafting HSC (LT-HSC) sub-population has been highly challenging (Dever et al., 2016; DeWitt et al., 2016; Genovese et al., 2014b; Hoban et al., 2015; Wang et al., 2015).

Cell cycle plays an important role in DNA repair decisions in response to double strand breaks. In most human cell types, NHEJ is the primary repair mechanism throughout the cell cycle, while HDR primarily occurs in S/G2 phase due to template availability and to avoid inappropriate telomere fusion during mitosis (Branzei and Foiani, 2008; Essers et al., 2002; Hustedt and Durocher, 2017; Mao et al., 2008; Orthwein et al., 2015; Pietras et al., 2011; Saleh-Gohari and Helleday, 2004). HSCs, like many other adult stem cells, can exist in both cycling and G0 quiescent states. For HSCs, cycling supports hematopoiesis, while quiescence preserves the stem population (Li and Clevers, 2010).

Quiescent HSCs from mice primarily employ NHEJ repair, while cycling mouse HSCs can employ both NHEJ and HDR (Beerman et al., 2014; Mohrin et al., 2010). But human HSCs are distinct from their mouse counterparts in terms of frequency of cycling (Abkowitz et al., 1996; Cheshier et al., 2007; Kiel et al., 2007), DNA damage response (Biechonski and Milyavsky, 2013; Mohrin et al., 2010), and expression of DSB repair genes (Biechonski and Milyavsky, 2013). The link between cell cycle status and the gene editing outcomes in human HSCs has yet been explored. Likewise, the use of single stranded oligonucleotide donors (ssODNs) during HSC editing shows therapeutic promise but is relatively unexplored from a mechanistic point of view (DeWitt et al., 2016; De Ravin et al., 2017).

Here we investigate the relationship between the cell cycle status of adult human mobilized peripheral blood (mPB) CD34+ HSPC subpopulations and their editing outcomes. We find that editing CD34+ HSPCs results in high levels of HDR in relatively differentiated subpopulations, but G0 HSPCs almost completely lack HDR alleles. Allowing HSPCs to briefly enter the cell cycle yields immunophenotypically primitive cells (CD34+ CD38-) with high levels of HDR but few quiescent cells. We define these CD34+ CD38-immunophenotypically primitive cells as “engraftment-enriched” (EE) HSPCs for the purpose of this paper since CD34+ CD38- HSPCs have been shown to primarily consist of cells that preserve the potential to engraft (Masiuk et al., 2017; Zonari et al., 2017). Using the timed administration of a small molecule cocktail originally developed for HSC maintenance, we developed a protocol to place HDR-edited EE HSPCs back into quiescence. The end result is G0 EE HSPCs whose HDR editing efficiency reflects the rest of the CD34+ HSPC population. This translates to an almost 6-fold increase in HDR/NHEJ ratios of EE HSPCs. These data yield new insights into the DNA repair preferences of HSPCs enriched for engrafting cells and suggests routes to therapeutic protocols for efficient genome editing to cure blood disorders.

## Results

### HSCs require more time to activate HDR pathways during gene editing as compared to differentiated cells

While gene editing reagents have been used to induce significant levels of HDR editing in bulk CD34+ HSPCs, maintenance of HDR for a prolonged period of time after *in vivo* engraftment has been challenging (Dever et al., 2016; DeWitt et al., 2016; Genovese et al., 2014b; Hoban et al., 2015; Wang et al., 2015). By contrast, NHEJ is maintained at high levels during prolonged engraftment. This could either arise because the act of editing somehow makes LT-HSCs lose markers of stemness, or because LT-HSCs do not perform HDR. To address this dichotomy, we first interrogated the extent to which primitiveness affects the repair decision after a Cas9-induced DSB in human mPB CD34+ HSPCs.

We utilized a potent single guide RNA (sgRNA) we previously found to efficiently edit human CD34+ HSPCs at the HBB locus and an ssODN donor template designed to modify the causative hemoglobin beta (*HBB*) mutation involved in sickle cell disease (SCD) (Figure S1A) (Cradick et al., 2013; DeWitt et al., 2016). After editing bulk CD34+ HSPCs, we measured the efficiency of HDR and NHEJ in immunophenotypically sorted HSCs (CD34+ CD38- CD45RA- CD90+), MPPs (CD34+ CD38- CD45RA- CD90-), and progenitors (CD34+ CD38+) (Figures 1A and 1B). Editing efficiency was quantified by using next-generation amplicon sequencing encompassing the HBB target site (Figure S1B).

We cultured CD34+ HSPCs in stem cell expansion media consisting of SFEMII and CC110 cytokine cocktail (SC) for one day, electroporated the cells with HBB-targeting Cas9 ribonucleoprotein complexes (RNPs) and cultured the HSPCs for one day before separating several HSPC subsets using fluorescence-activated cell sorting (FACS) and assessing the editing efficiency in each subset through NGS genotyping (Figure 1C top). Both HDR and NHEJ were evident in bulk CD34+ cells and relatively differentiated progenitors (CD34+ CD38+). Total editing was somewhat reduced in MPPs (CD34+ CD38- CD45RA- CD90-). Strikingly, we found moderate amounts of NHEJ in immunophenotypic HSCs (CD34+ CD38- CD45RA- CD90+) but almost no HDR in these cells, which led to a 3-fold lower HDR/NHEJ ratio in HSCs compared to bulk CD34+ HSPCs. (Figure 1C).

We further cultured the sorted populations (HSCs, MPPs, and progenitors) and found that HSCs eventually accumulated HDR edits, but only 72 hours after electroporation (Figure S1C). However, the HDR/NHEJ ratio was highest in progenitors and lowest in HSCs even 72 hours after electroporation (Figure S1C). In contrast, keeping CD34+ HSPCs in culture for two days before electroporation led to the appearance of significant HDR edits just one day after electroporation (Figure 1D). HDR was evident in all HSPC subtypes, including HSCs. These data indicate that more primitive HSCs preferentially repair Cas9-induced DSBs via NHEJ, but additional time in culture prior to the introduction of a DSB activates pathways related to HDR.

### Establishing the timing of cell cycle status in CD34+ subsets during *ex vivo* culture

HSPC primitiveness is linked to slower entry into the cell cycle (Laurenti et al., 2015) as well as lower frequency of cell cycle (Bradford et al., 1997; Morrison and Weissman, 1994; Pietrzyk et al., 1985; Suda et al., 1983; Uchida et al., 2003), and cell cycle progression is a major hallmark of increasing time in culture for HSPCs. Since HDR is intimately linked with cell cycle, we hypothesized that HSCs cannot utilize HDR at short culture time points due to quiescence resulting from slow entry into the cell cycle.

While the cycling properties of freshly isolated mouse and human HSC subpopulations have been described (Benveniste et al., 2010; Cheshier et al., 1999; Copley et al., 2012; Foudi et al., 2009; Laurenti et al., 2015; Oguro et al., 2013; Passegué et al., 2005; Qiu et al., 2014; Wilson et al., 2008), the cycling properties of human CD34+ HSPCs during extended *ex vivo* culture are not fully established. Before investigating the relationship between cell cycle status and editing efficiency, we first explored the cell cycle progression of CD34+ cells in *ex vivo* culture using immunophenotyping combined with Hoechst 33342 (stains for DNA) and Ki67 (highly expressed in proliferating cells) staining. (Gerdes et al., 1984; Kim and Sederstrom, 2015). We found that more than 50% of cryopreserved mPB CD34+ HSPCs are quiescent (in G0) when thawed, but they gradually enter the cell cycle and are fully cycling by 3 days in SC culture (Figures 2A, 2B, and S2).

**Figure 2.**
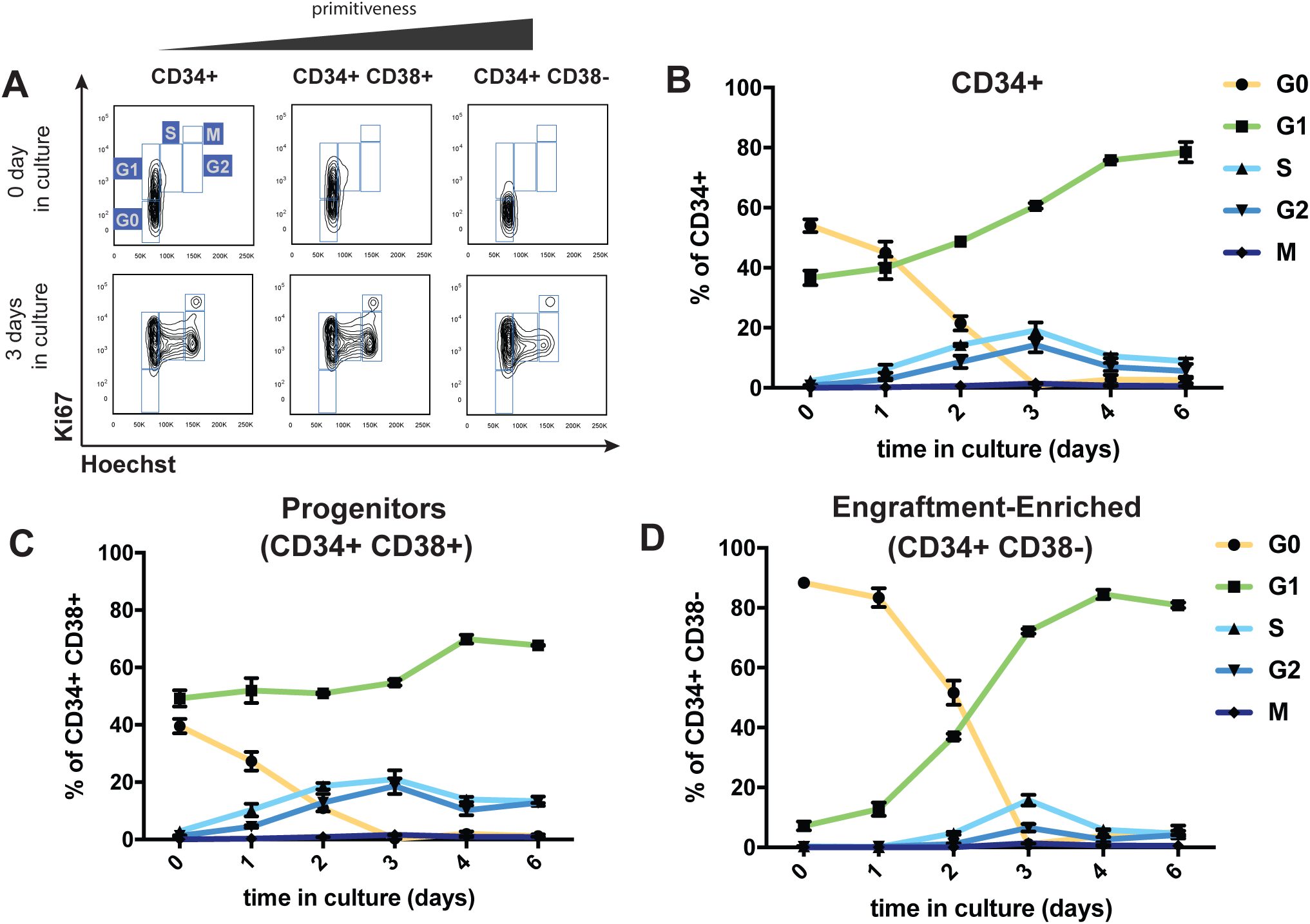
Cell cycle progression of human mPB CD34+ cells in *ex vivo* culture. (A) Representative flow cytometry plots for assessing cell cycle status in CD34+, CD34+ CD38+ (Progenitors), and CD34+ CD38- (Engraftment-enriched) populations. CD34+ cells were stained with antibodies against Ki67 (highly expressed in proliferating cells) and Hoechst 33342 (stains for DNA). G0: 2N DNA and Ki67 negative, G1: 2N DNA and Ki67 positive, S/G2/M: 4N DNA and Ki67 positive. (B) Cell cycle status of CD34+ cells in *ex vivo* culture. ~60% of bulk CD34+ cells are in G0 at day 0 and 100% of the cells are cycling by day 3. (C) Cell cycle status of CD34+38+ progenitor cells in *ex vivo* culture. More than 50% of progenitor cells are cycling at day 0 and the percentage of cycling cells continually increase until day 3. (D) Cell cycle status of CD34+38- engraftment-enriched (EE) HSPCs in *ex vivo* culture. ~90% of the EE HSPCs are quiescent at day 0 and EE HSPCs gradually exit quiescence until day 3 where 100% of them are cycling. Representative data from n≥2 independent experiments with n=2 biological replicates for each condition. *See also Figure S2*.

We next examined CD34+ CD38- HSPCs, which contain most of the engraftment potential within the CD34+ population and are enriched for primitive populations such as HSCs and MPPs, as well as the more differentiated progenitor CD34+ CD38+ populations. For simplicity, here we define CD34+ CD38- HSPCs as “engraftment-enriched” (EE) HSPCs. Interestingly, EE HSPCs have a delayed exit from quiescence compared to progenitors (Figures 2A, 2C, 2D, and S2). When cells are edited after only one day in culture (Figure 1C), 80% of EE HSPCs are quiescent at the time of editing while 70% of CD34+ CD38+ progenitors are cycling (Figures 2C, 2D, and S2). These results support the absence of HDR in quiescent cells and correlate with the vast difference in HDR efficiency between HSCs and progenitors (Figure 1C). By contrast, when cells are edited after two days in culture (Figure 1D), more than 50% of the EE HSPCs have begun to actively cycle. This could account for the significant amount of HDR observed in HSCs during longer *ex vivo* culture (Figures 1D, 2C, and 2D).

### Quiescent CD34+ HSPCs perform only NHEJ but cycling CD34+ HSPCs perform both NHEJ and HDR

To directly test how cell cycle status affects editing of human adult stem cells, we edited CD34+ HSPCs after one day in culture, allowed them to resolve edits for another day in culture, sorted them by cell cycle status, and used amplicon NGS to assess each population’s editing outcomes (Figure 3A top, Figure S3A). One day after editing we found that cells in G1 and S- G2-M stages had a substantial amount of HDR alleles, but quiescent G0 CD34+ HSPCs almost completely lacked HDR alleles and had 3-fold decrease in HDR/NHEJ ratio compared to cycling HSPCs (Figure 3A). We observed NHEJ alleles in significant amounts regardless of cell cycle, though the highest amount was observed in the S-G2-M population (Figure 3A). Intriguingly, six hours after editing we found small amounts of NHEJ alleles across various cell cycle subpopulations, but HDR alleles do not appear in any of the cell cycle subpopulations, consistent with reports from other cell types that HDR takes longer than NHEJ (Arnoult et al., 2017; Mao et al., 2008) (Figure S3B).

**Figure 3.**
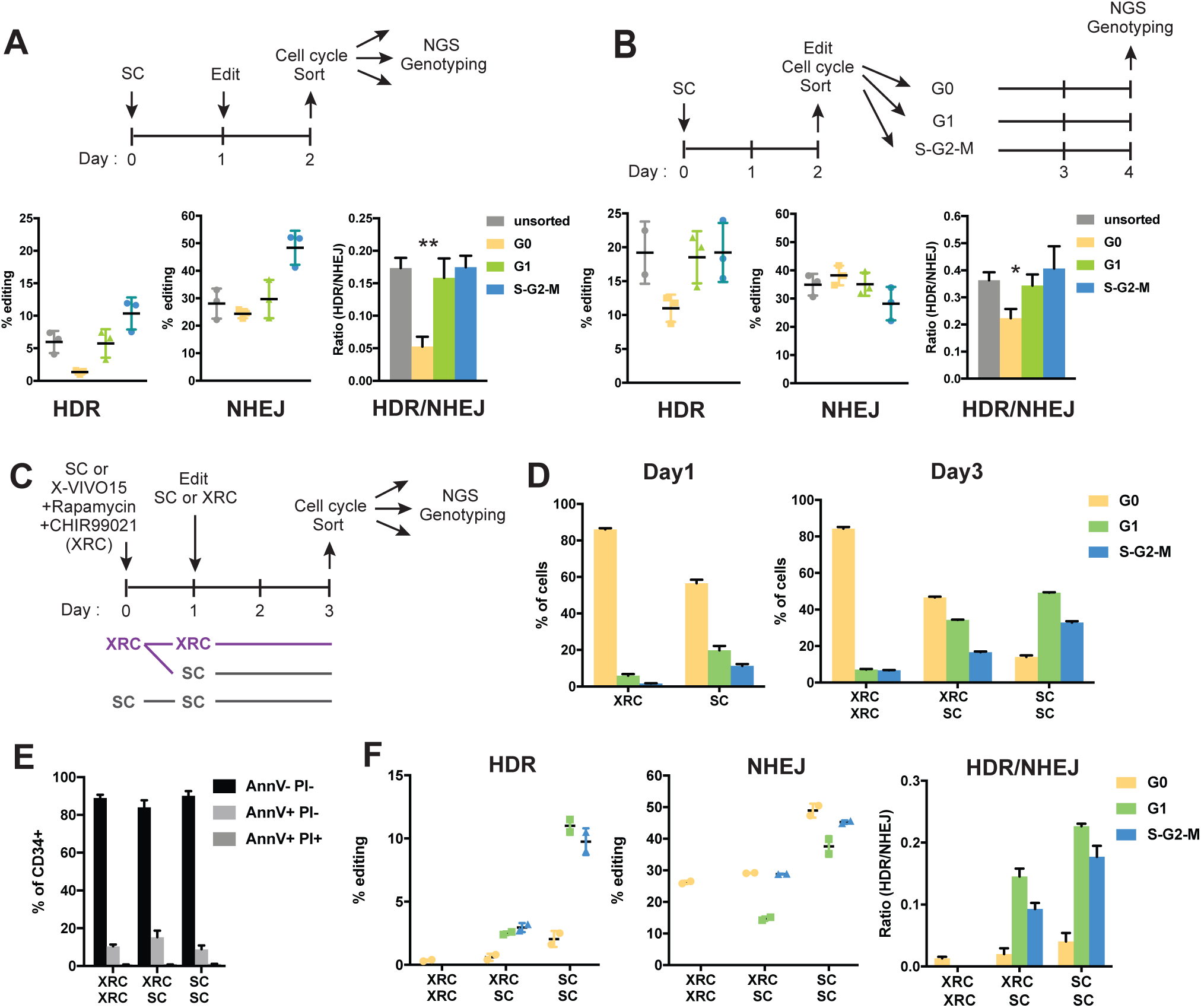
Quiescent CD34+ HSPCs perform only NHEJ, but cycling CD34+ HSPCs perform both NHEJ and HDR. (A) Editing outcomes in CD34+ subpopulations in different cell cycle status. 1 day post electroporation, 2 days in culture. Percentage of reads positive for HDR or NHEJ by next-generation amplicon sequencing at the HBB site. G0 CD34+ HSPCs in 2-day culture do not accumulate HDR alleles. Representative data from n≥2 independent experiments with n≥2 biological replicates for each condition. Mean ± SD shown. **: p<0.01 by unpaired t-test. (B) Editing outcomes in CD34+ subpopulations sorted into different cell cycle statuses. 2 days post electroporation, 4 days in culture. Percentage of reads positive for HDR or NHEJ by next-generation amplicon sequencing at the HBB site. G0 CD34+ HSPCs sorted at 2 days in culture accumulates significant HDR alleles in 4-day culture. Representative data from n≥2 independent experiments with n≥2 biological replicates for each condition. Mean ± SD shown. *: p<0.05 by unpaired t-test. (C) Schematic of the workflow for mTOR and GSK-3 inhibition with rapamycin and CHIR99021 (XRC) for inhibition of cell cycle entry. Culture condition: SFEMII + CC110 (SC), X-VIVO15 + Rapamycin + CHIR99021 (XRC) (D) Cell cycle profiles of CD34+ cells in SC or XRC media at the time of electroporation (Day1) and two days post nucleofection (Day3). XRC media prevents cell cycle entry but this can be reversed by placing the CD34+ cells in SC media. n=2 biological replicates for each condition. Mean ± SD shown. (E) Percentage of early and late apoptosis was assessed by staining the cells for Annexin V (AnnV) and Propidium Iodide(PI) at 2 days post nucleofection. AnnV-PI-: viable; AnnV+PI-:early apoptotic; AnnV+PI+; apoptotic. XRC media does not induce apoptosis. n=2 biological replicates for each condition. Mean ± SD shown. (F) Editing outcomes in CD34+ cells kept in SC or XRC media in different cell cycle status 2 days post nucleofection, 3 days in culture. Inhibition of cell cycle entry by XRC blocks HDR repair but is reversible. n=2 biological replicates for each condition. Mean ± SD shown. *See also Figure S3*.

We next asked whether additional time in culture altered CD34+ HSPC editing outcomes according to cell cycle status. Since Hoechst staining led to a significant decrease in viability in CD34+ HSPCs (Figure S3C and S3D), we developed a live cell staining protocol that utilizes Pyronin Y that can stain both DNA and RNA when used alone (Darzynkiewicz et al., 1987, 2004). Cells were cultured for two days, then edited and immediately subjected to a live cell cycle sort using Pyronin Y accumulation (Figure 3B top and Figure S3E). Sorted subpopulations were cultured for an additional 2 days before NGS genotyping in order to allow edits to resolve according to cell cycle status (Figure 3B top). Similar to short-culture experiments, we observed NHEJ in cells regardless of cell cycle (Figure 3B middle). Unlike short-culture experiments, quiescent G0 cells kept in culture for a total of four days displayed substantial HDR alleles (Figure 3B left).

Since almost all CD34+ cells exit quiescence within three days in culture (Figure 2), CD34+ HSPCs that are still in G0 at the time of editing (mostly CD34+ CD38- EE HSPCs) should exit quiescence by the end of a long culture and would be able to accumulate significant HDR alleles while cycling. Hence, our results overall suggest that non-cycling CD34+ HSPCs in G0 are highly enriched in primitive EE HSPCs and heavily rely on the NHEJ pathway as opposed to HDR. By contrast, cycling cells in G1 and S-G2-M are enriched in more differentiated CD34+ CD38+ progenitors and utilize both HDR and NHEJ.

### Preventing exit from quiescence blocks HDR repair in CD34+ HSPCs

Our previous experiments showed that quiescent, primitive HSPC subsets are less likely to perform HDR than cycling, differentiated subsets. We next tested whether induction of quiescence was sufficient to affect HDR levels under otherwise HDR-competent conditions.

We induced quiescence using either retinoic acid, which has been shown to drive mouse HSCs into deep dormancy (Cabezas-Wallscheid et al., 2017), or inhibitors of mTOR (Rapamycin) and GSK-3 (CHIR9901), which have been used to maintain mouse and human HSCs *ex vivo* and *in vivo* (Huang et al., 2012) (Figure S3F). We found that treatment of CD34+ HSPCs with retinoic acid in SC media led to differentiation as measured by substantial loss of CD34 expression which could potentially be due to the differences in maintenance of HSCs in mouse and human, whereas a combination of Rapamycin and CHIR99021 in X-VIVO media (XRC) led to the prevention of cell cycle entry while maintaining primitiveness (Figure S3F-H).

We investigated editing outcomes in CD34+ HSPCs cultured in XRC media as compared to SC expansion media. We used three different treatment regimens (Figure 3C). One set of HSPCs was kept in SC both before and after editing. A second set was started in XRC before editing, and then either maintained in XRC after editing or moved to SC after editing. All cells were sorted based on cell cycle and editing outcomes for each stage of the cell cycle were measured by NGS. Cells maintained in SC media entered cell cycle as normal, exhibiting a decrease in G0 cells and increase in G1 and S-G2-M cells after three days. Pretreatment of CD34+ HSPCs with XRC media led to the prevention of cell cycle entry, with almost all cells in G0 after three days (Figure 3D). Treatment with XRC was not associated with a decrease in cell viability (Figure 3E), and moving XRC-treated cells to SC media allowed HSPCs to re-enter the cell cycle, as measured by a decrease in G0 cells and increase in G1 and S-G2-M. (Figure 3D).

Strikingly, quiescent CD34+ HSPCs treated continuously with XRC repaired almost all Cas9-induced DSBs using NHEJ and harbored almost undetectable levels of HDR alleles. Moving XRC-treated HSPCs to SC media after editing led to increased levels of HDR, but this was mostly confined to cells in G1 and S-G2-M. HSPCs maintained in SC before and after editing exhibited low levels of HDR in G0 cells, but high levels of HDR in G1 and S-G2-M (Figures 3F). These results show that small molecule-induced quiescence in HSPCs is sufficient to prevent HDR even after multiple days in *ex vivo* culture, and that cycling is necessary for high levels of HDR.

### Inducing quiescence after a short period of cycling yields quiescent, primitive HSPCs that harbor HDR alleles

While XRC treatment has previously been used to maintain stemness (Huang et al., 2012), we next asked whether these compounds could induce quiescence after HSPCs have been allowed to cycle. Our overall goal was to allow HSPCs to cycle to accumulate HDR alleles during editing, and then to place them back into G0 in order to maintain stemness.

We edited CD34+ HSPCs and cultured them in SC media to allow them to enter the cell cycle (Figure 4A). On the day of electroporation, 50% of CD34+ HSPCs were quiescent as expected (Figure S4A). Two days after editing we sorted cells based on cell cycle and quantified editing outcomes by NGS. We found that at this timepoint most cells had exited G0 and were in G1 or S-G2-M (Figure 4B) although less of EE HSPCs were in S-G2-M compared to the progenitors (Figure S4B). As before, HDR alleles were almost completely absent from the remaining G0 cells but present in G1 and S-G2-M cells, while NHEJ alleles were present in all stages of the cell cycle (Figure 4C). HDR/NHEJ ratio was 7 times lower in G0 cells than G1 and S-G2-M cells. We then kept the remaining HSPCs in SC to allow cycling to continue for another three days, or moved them to XRC to induce quiescence. Six days after editing (three days in SC and three additional days in either SC or XRC), we sorted based on cell cycle and quantified repair outcomes by NGS.

**Figure 4.**
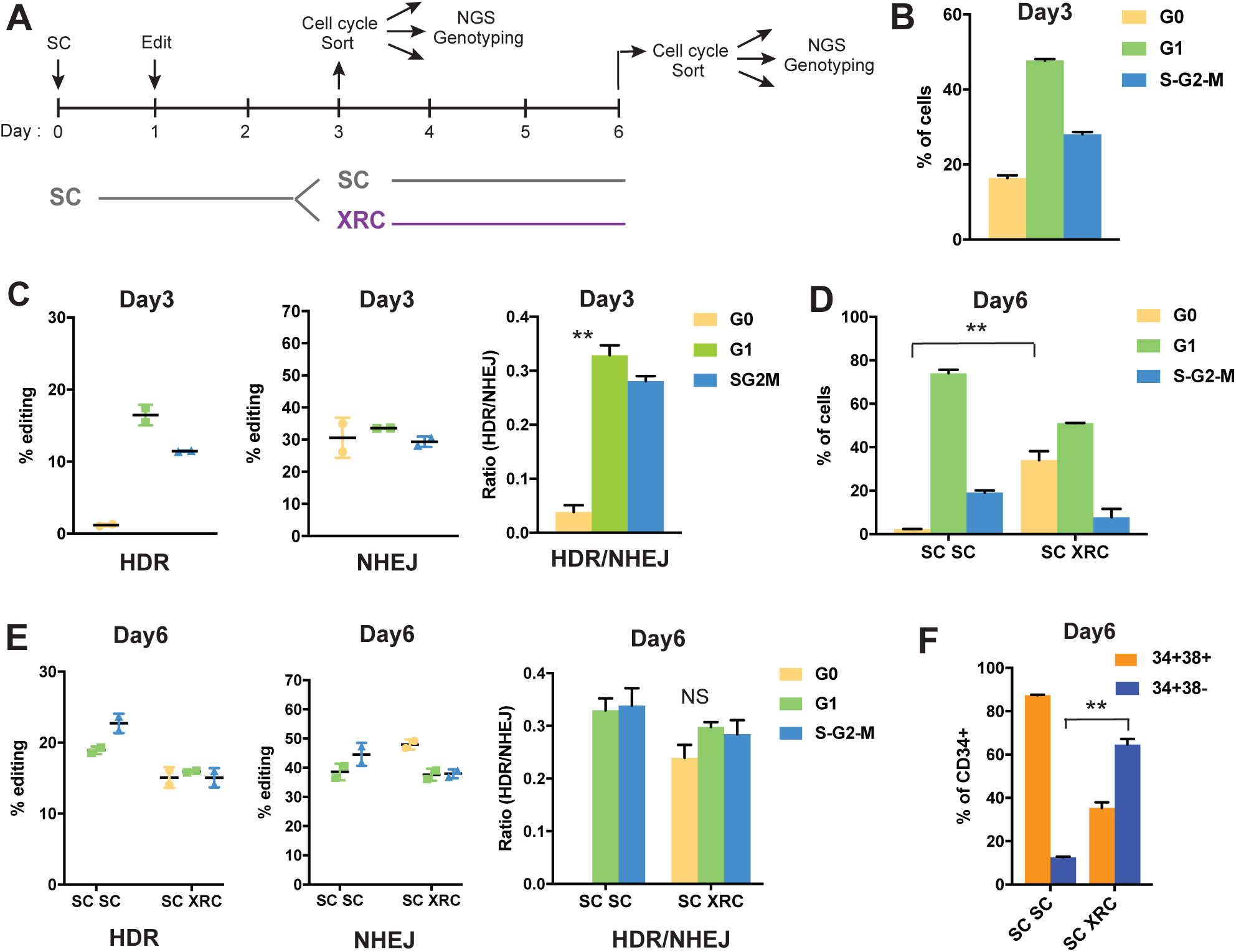
Inducing quiescence after a short period of cycling yields quiescent, primitive HSPCs that harbor HDR alleles. (A) Schematic of the workflow for inducing quiescence after a short period of cycling. CD34+ HSPCs are placed in SC culture for 1 day before editing, cycles for 2 additional days in SC culture, then quiescence is induced with XRC for 3 days before the cells are subjected to FACS based on their cell cycle status and genotyped by NGS. (B) Cell cycle profiles of CD34+ cells 3 days in culture (SC). Majority of CD34+ HSPCs are cycling at day 3. Representative data from n≥3 independent experiments with n≥2 biological replicates for each condition. Mean ± SD shown. (C) Editing outcomes in CD34+ cells 3 days in culture (SC). HDR repair does not take place in G0 CD34+ HSPCs. Representative data from n≥3 independent experiments with n≥2 biological replicates for each condition. Mean ± SD shown. **: p<0.01 by unpaired t-test. (D) Cell cycle profiles of CD34+ cells 6 days in culture (3 days in SC media and 3 additional days in SC or XRC media). 3 days in XRC media induces quiescence in 30% of the cells. Representative data from n≥3 independent experiments with n≥2 biological replicates for each condition. Mean ± SD shown. **: p<0.01 by unpaired t-test. (E) Editing outcomes in CD34+ cells 6 days in culture (3 days in SC media and 3 additional days in SC or XRC media). 3 days in XRC media results in HDR edits in G0 CD34+ HSPCs. Representative data from n≥3 independent experiments with n≥2 biological replicates for each condition. Mean ± SD shown. (F) Percentage of CD34+ cells that are CD34+CD38+ (progenitors) vs. CD34+CD38- (EE HSPCs) 6 days in culture. CD34+ cells that regained quiescence in XRC media includes higher proportion of EE HSPCs compared to CD34+ cells maintained soley in SC media. n=2 biological replicates for each condition. Mean ± SD shown. **: p<0.01 by unpaired t-test. *See also Figure S4*.

HSPCs that were maintained continuously in SC media were almost completely lacking in G0 cells by six days after editing (Figure 4D). The remaining cells, which were only in G1 or S-G2-M, harbored high levels of HDR alleles (Figure 4E). By contrast, cells moved to XRC media had almost 40% G0 cells and relatively few cells in S-G2-M (Figure 4D). Notably, 60% of the primitive EE HSPCs returned to quiescence as compared to 30% of CD34+38+ progenitors (Figure S4E). We found that the G0 cells in XRC now harbored high levels of HDR alleles, and were in fact comparable in HDR to G1 and S-G2-M cells (Figure 4E). We show that XRC treatment maintains stemness (Figure S4C) and supports viability (Figure S4D) and is distinctive from the omission of cytokines that generally leads to loss of CD34+ expression and viability (Figure S4C and S4D). We further found that post-treatment with XRC led to enrichment in EE HSPCs (CD34+ CD38- markers of stemness) as compared to continued culture in SC (Figure 4F). In sum, we have developed a strategy to enable high efficiency HDR in immunophenotypically primitive and quiescent HSPCs. This strategy allows HSPCs to briefly cycle after Cas9-mediated induction of a DSB to allow HDR, and then places cells back into quiescence after HDR alleles have been acquired.

## Discussion

Our data suggest a potential path towards the use of gene edited HSCs for therapeutic purposes, and also shed light on fundamental HSC biology. The DNA repair decisions after a double strand break in primitive human hematopoietic cells are essential for cell survival, yet are underexplored due to difficulties in studying human HSCs. Aged human hematopoietic cells shows elevated levels of unresolved DSBs and increased mutation frequencies, but the mechanisms underlying DSB repair in these cell types are largely unknown (Beerman et al., 2014; Genovese et al., 2014a; Rossi et al., 2007; Rübe et al., 2011; Weinstein et al., 2013). Here we have used CRISPR genome editing to induce a precise DSB in mixed CD34+ HSPCs and measured its repair in various cell subtypes and phases of the cell cycle via a combination of FACS and NGS. This approach could broadly accelerate in-depth probing of DNA repair decisions in many different stem cell types.

We found that genome editing of CD34+ HSPCs leads to high levels of NHEJ in multiple cell subtypes, but that HDR is preferentially missing from more primitive quiescent cells. Instead, HDR accumulates in relatively differentiated cells and immunophenotypically primitive cells that have exited quiescence. Several groups have reported that genome editing CD34+ HSPCs leads to high-efficiency HDR in relatively short term *in vitro* culture that drops dramatically during long term *in vivo* engraftment (Dever et al., 2016; DeWitt et al., 2016; Genovese et al., 2014b; Hoban et al., 2015; Wang et al., 2015). This is true even with very different modalities of Cas9 (mRNA, recombinant protein), guide RNA (synthetic, AAV-expressed), and HDR donor (single stranded DNA, AAV6) (Dever et al., 2016; DeWitt et al., 2016; Genovese et al., 2014b; Hoban et al., 2015; Kim et al., 2014; Wang et al., 2015). Our results suggest that the observed *in vivo* lack of HDR is caused not by an inability to target immunophenotypic LT-HSCs, but because the repopulating stem cells are in an inappropriate phase of the cell cycle to perform HDR.

Mechanistic investigations of DNA repair have established that HDR is preferentially active in the S/G2 stages of the cell cycle, probably to avoid deleterious telomere fusions that can occur if HDR is active during mitosis (Orthwein et al., 2014). Stem cells, such as HSCs therefore represent a challenge in that they divide during self-renewal and differentiation, but their stemness is intricately linked to long-term quiescence (Ema et al., 2000; Morrison and Weissman, 1994; Suda et al., 1983). Prolonged *in vitro* culture of HSCs leads to a loss of stemness, entry into the cell cycle, and poor engraftment (Morrison and Kimble, 2006; Wilson et al., 2008). There is a therefore a fundamental tension between HDR editing and the maintenance of stemness via quiescence. LT-HSCs may need to lose a defining feature of their stemness in order to obtain HDR edits.

One might avoid HDR entirely and instead pursue NHEJ-based editing. This approach shows promise for the treatment of sickle cell disease, where disruption of various repressor elements leads to re-expression of protective fetal hemoglobin (Bauer et al., 2013; Bjurström et al., 2016; Canver et al., 2015; Chang et al., 2017). NHEJ is well-represented in long-term engrafting HSCs during genome editing (Dever et al., 2016; DeWitt et al., 2016; Genovese et al., 2014b; Hoban et al., 2015; Wang et al., 2015), and here we show via immunophenotyping and cell cycle analysis that HSCs in G0 are fully capable of accumulating NHEJ alleles. However, limiting oneself to NHEJ-based editing does not fully tap the potential of genome editing. Many fundamental questions are best answered by surgically replacing genomic sequences, and relatively few genetic diseases can be cured by NHEJ-based sequence disruption.

We find that the drop in levels of HDR after long term CD34+ engraftment is reflected in poor HDR in the quiescent LT-HSC subpopulation. By contrast, cycling progenitor cells and even MPPs exhibit significant levels of HDR. Since HDR is maximal during S-G2 phase, we reasoned that progression through at least one cell cycle would be required for efficient HDR in HSCs (Branzei and Foiani, 2008; Hustedt and Durocher, 2017). We show that allowing quiescent EE HSPCs to briefly enter the cell cycle enables HDR while retaining immunophenotypic markers representative of engraftment potential. We therefore allowed HSPCs to exit G0, edited them, allowed them to cycle, and only later added XRC to induce quiescence. This strategy results in quiescent EE HSPCs that exhibit more than 6-fold increases in HDR, close to those observed in cycling progenitors.

Directly addressing the tension between quiescent stemness and HDR is critical to fully achieve the potential of genome editing. Multiple types of stem cells potentially suffer from poor HDR that may be linked to quiescence (Bressan et al., 2017; Schwank et al., 2013; Urnov et al., 2010; Zhu et al., 2017). It remains to be seen if the re-quiescence strategy we describe here will be applicable beyond HSCs, though one barrier is the paucity of culture models for various types of stem cells. Our data indicate that culture conditions can be just as important as editing modality to achieve desired genomic outcomes.

## Acknowledgments

J.E.C., S.K.W., and N.L.B. are supported by the Li Ka Shing Foundation and Heritage Medical Research Institute. J.E.C. is supported by CIRM under TRAN1-09292 and the National Heart, Lung, and Blood Insitute of the NIH under DP2-HL-141006. J.S. is supported by the National Institute on Aging of the NIH under T32 AG000266. M.A.D. and J.V. are supported by CIRM TRAN1-09292.

## Author Contributions

J.S. and J.E.C. conceived and designed the experiments. J.S., M.A.D., and J.V. carried out experiments. S.K.W. and N.L.B. carried out bioinformatics analysis. J.S. and J.E.C. wrote the paper.

## STAR Methods

### CONTACT FOR REAGENT AND RESOURCE SHARING

Further information and requests for reagents may be directed to, and will be fulfilled by, the Lead Contact

### EXPERIMENTAL MODEL AND SUBJECT DETAILS

Cryopreserved wildtype human mobilized peripheral blood CD34+ HSPCs from multiple volunteer donors including male and female whose age ranged from 20-35 were purchased from Allcells, Inc.

### METHOD DETAILS

#### Primary Cell Culture

CD34+ HSPCs were cultured in SC (SFEMII + CC110 (StemCell Technologies)) media, XRC (X-VIVO15 (Lonza) + 5nM Rapamycin (EMD Millipore) + 3uM CHIR99021 (EMD Millipore)), or SC + 5uM All-trans retinoic acid (Sigma) media unless otherwise noted.

#### Electroporation for editing experiments

Cas9 RNP synthesis was carried out as previously described (Lin, Dewitt). Briefly, 75pmol of Cas9-NLS (UC Berkeley, Berkeley, CA) was mixed slowly into Cas9 buffer (20mM HEPES (pH 7.5), 150mM KCl, 1mM MgCl_2_, 10% glycerol and 1mM TCEP) containing 75pmol of synthetic sgRNA targeting the HBB locus (Synthego). The resulting 7.5ul mixture was incubated for 15minutes to allow RNP formation. 2x10^-5^ CD34+ HSPCs were harvested, washed once with PBS, and resuspended in 20ul of P3 nucleofection buffer (Lonza, Basel, Switzerland). 7.5ul of RNP mixture and 20ul of cell suspension were combined and added into a Lonza 4d strip nucleocuvette and were electroporated with program ER-100. 200ul pre-warmed media was added to each nucleocuvette and electroporated cells were transferred to culture dishes. Editing outcomes were measured 1-5 days post-electroporation by Next Generation Amplicon Sequencing.

#### PCR and Next-Generation Amplicon Sequencing preparation

50-100ng of genomic DNA from edited CD34+ cells was amplified at HBB sites using primer set 1 (Figure S1A). The PCR products were SPRI cleaned, followed by amplification of 20-50ng of the first PCR product in a second 12 cycle PCR using primer set 2 (Figure S1A). Then the second PCR products were SPRI cleaned, followed by amplification of 20-50ng of the second PCR product in a third 9 cycle PCR using illlumina compatible primers (primers designed and purchased through the Vincent J. Coates Genomics Sequencing Laboratory (GSL) at University of California, Berkeley), generating indexed amplicons of an appropriate length for NGS. Libraries from 100-500 pools of edited cells were pooled and submitted to the GSL for paired-end 300 cycle processing using a version 3 Illumina MiSeq sequencing kit (Illumina Inc., San Diego, CA) after quantitative PCR measurement to determine molarity.

#### Next-Generation Amplicon Sequencing analysis

Samples were deep sequenced on an Illumina MiSeq at 300bp paired-end reads to a depth of at least 10,000 reads. A modified version of CRISPResso (Pinello et al., 2016) was used to analyze editing outcomes. Briefly, reads were adapter trimmed then joined before performing a global alignment between reads and the reference and donor sequences using NEEDLE (Li et al., 2015). Rates of HDR are calculated as total reads that successfully convert the main (non-PAM out) edit site and have no insertions or deletions within three basepairs to each side of the cutsite divided by the total number of reads. NHEJ rates are calculated as any reads where an insertion or deletion overlaps the cutsite or occurs within three basepairs of either side of the cutsite divided by the total number of reads.

#### Immunofluorescence

##### Immunophenotypic analysis assays

Human CD34+ cells with or without editing were were first stained with fixable viability stain 660 (1:1000, BD) for 5 min in 37°C and then were stained with Percp-Cy5.5-anti-CD34 (1:50), PE-Cy7-anti-CD38 (1:50), PE-anti-CD90 (1:30), FITC-anti-CD45RA (1:25), and BV421-anti-CD49f (1:30) (all of the antibodies are from BD) for 30 min in 4**°**C. Samples were then sorted on Aria Fusion Cell Sorter (BD) or analyzed on LSR Fortessa cytometer (BD).

##### Cell cycle analysis assays

For Ki67-Hoechst assays, CD34+ cells with or without editing were first stained with fixable viability stain 660 (1:1000, BD) for 5 min in 37°C and were fixed with Cytofix/Cytoperm buffer (BD) for 15 min in 4°C. Cells were stained with FITC-anti-KI67 (1:25, BD, 556027) for 2hours-overnight in Permwash buffer (BD), then with Hoechst 33342 (1:5000; Life Technologies) for 5 min at RT. Samples were sorted on a Aria Fusion Cell Sorter (BD) or analyzed on a LSR Fortessa cytometer (BD). For assessment of immunophenotypic markers together with cell cycle analysis, human CD34+ cells with or without editing were stained with Percp-Cy5.5-anti-CD34 (1:50) and PE-Cy7-anti-CD38 (1:50) for 30 min in 4°C before they were stained with fixable viability stain and fixed. For assessment of cell cycle status without fixation (live cell cycle status), cells with or without editing were stained with Hoechst 33342 (1:1000, Invitrogen) for 45min in 37°C, and then were stained with Pyronin Y (1:20,000, Invitrogen) for additional 15 min in 37°C or were just stained with Pyronin Y for 15 min in 37°C. Samples were then sorted on Aria Fusion Cell Sorter (BD) or analyzed on LSR Fortessa cytometer (BD).

##### Apoptosis analysis assays (Annexin V, PI)

Human CD34+ cells with or without editing were first stained with Percp-Cy5.5-anti-CD34 (1:50) and PE-Cy7-anti-CD38 (1:50) for 30 min in 4°C before they were washed twice with BioLegend’s Cell Staining Buffer (Biolegend) and stained with FITC Annexin V (Biolegend, 1:20) and PI (Biolegend, 1:10) for 15 minutes at room temperature. Then 400ul of Annexin V binding buffer was added and the samples were analyzed by LSR Fortessa cytometer (BD).

### QUANTIFICATION AND STATISTICAL ANALYSIS

Statistical analyses were performed with GraphPad Prism (version 7.00 for Mac, GraphPad Software) using unpaired two-tailed t-test analysis. Representative data from n≥2 independent experiments are shown in the figures unless otherwise stated. Each experiment included n≥2 biological replicates unless otherwise noted. More detailed information of experimental replicates are given in the figure legends of the corresponding experiments. All values are given in the text as mean (±SD) and a p value < 0.05 was accepted as significant in all analyses, unless otherwise stated.

### DATA AND SOFTWARE AVAILABILITY

The GEOS accession numbers for the next-generation sequencing data reported in this paper are ### (To be added).

#### Supplemental item titles and legends

**Figure S1.**
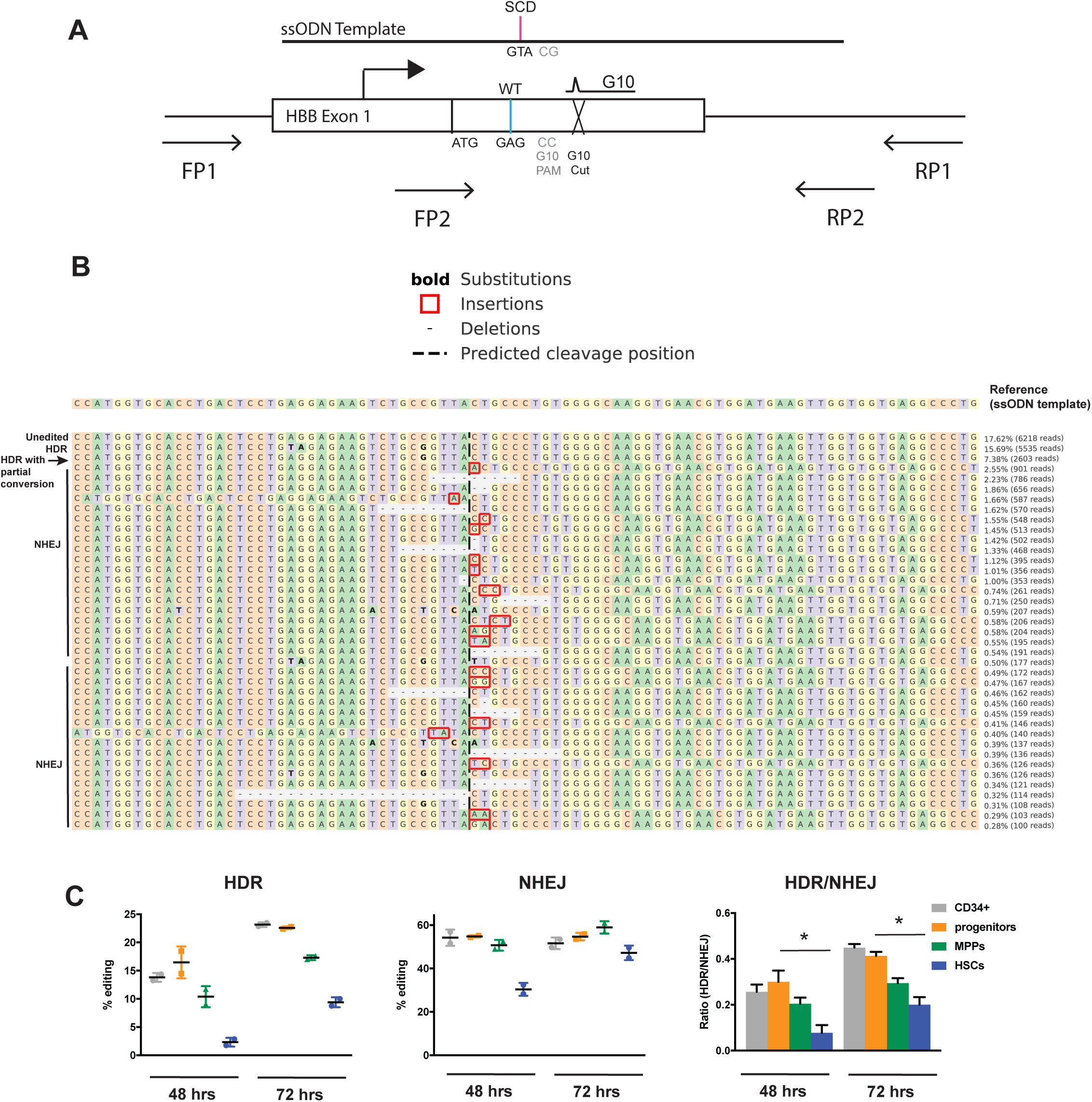
*Related to Figure 1*: Targeted gene editing at the HBB locus. (A) Schematic of the ssODN template, sgRNA (G10) designed to modify the causative hemoglobin beta (HBB) mutation involved in sickle cell diseasease (SCD), and PCR primers used for amplicon NGS library preparation to assess HDR and NHEJ efficiency. (FP1: tcacttagacctcaccctgtg, RP1: tatgggacgcttgatgttttct, FP2: tatgggacgcttgatgttttct, RP2: ctctgcctattggtctattttccca) (B) Example of an editing result from the amplicon NGS pipeline. (C) HBB target editing efficiency in CD34+ subpopulations after 48 and 72 hours of electroporation. Mean ± SD shown. *:p<0.05 by unpaired t-test.

**Figure S2.**
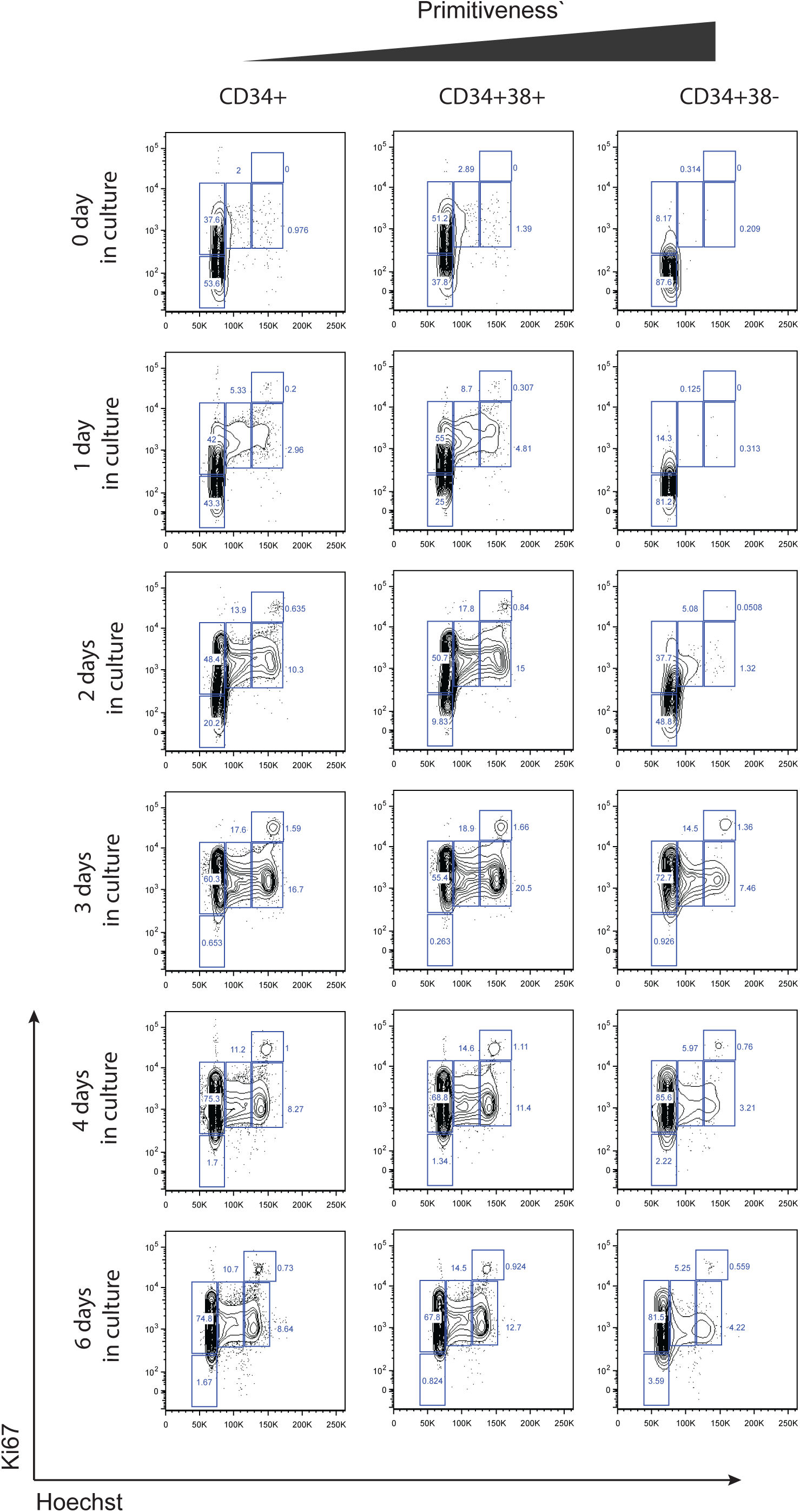
*Related to Figure 2*: Cell cycle progression of human mPB CD34+ HSPCs in ex vivo culture. Percentage of G0, G1, S, G2, and M cells in CD34+, CD34+ CD38+ progenitors, and CD34+ CD38- engraftment-enriched (EE) HSPCs after 0-6 days in ex vivo SC culture.

**Figure S3.**
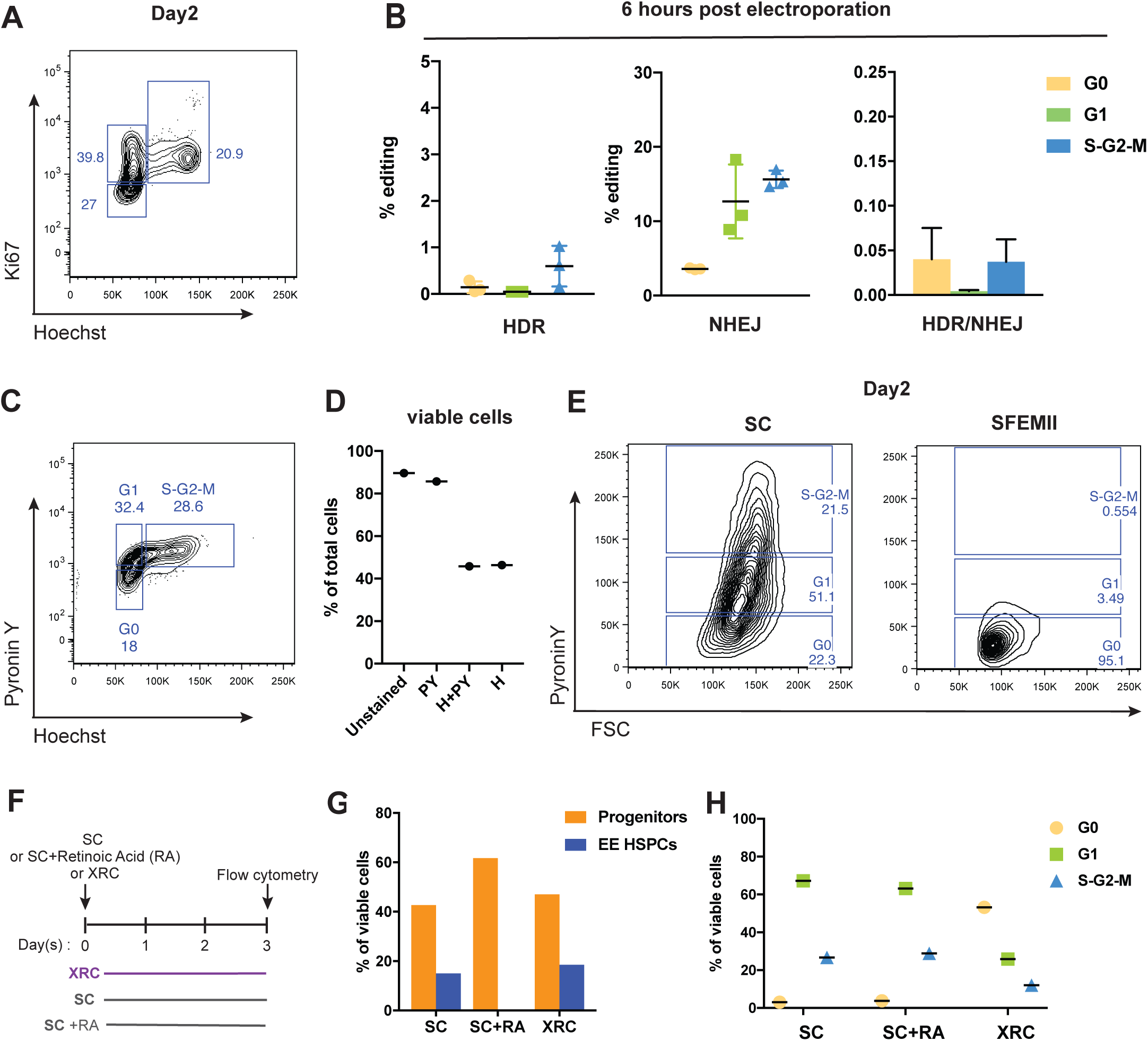
*Related to Figure 3*: Assessing editing efficiency in CD34+ HSPCs that are in different cell cycle status and establishing an ex vivo culture protocol that maintains quiescence and stemness of CD34+ HSPCs. (A) Representative flow plot for measuring cell cycle status in Figure 3A. (B) Editing outcomes in CD34+ population in different cell cycle status 6 hours after electroporation. Mean ± SD shown. (C) Representative flow plot for live cell cycle measurement. (D) Viability 2 days after staining with Pyronin Y, Hoechst 33342, or both. (E) Representative flow plot for measuring live cell cycle status using Pyronin Y in Figure 3B. (F) Schematic for the testing of ex vivo culture protocol (SC, SC + Retinoic Acid, XRC) to maintain quiescence without the loss of stemness measured by % of EE HSPCs. (G) Percentage of progenitors (CD34+ CD38+) vs. EE HSPCs (CD34+ CD38-) among viable cells. (H) Percentage of CD34+ HSPCs in G0, G1, and S-G2-M among viable cells.

**Figure S4.**
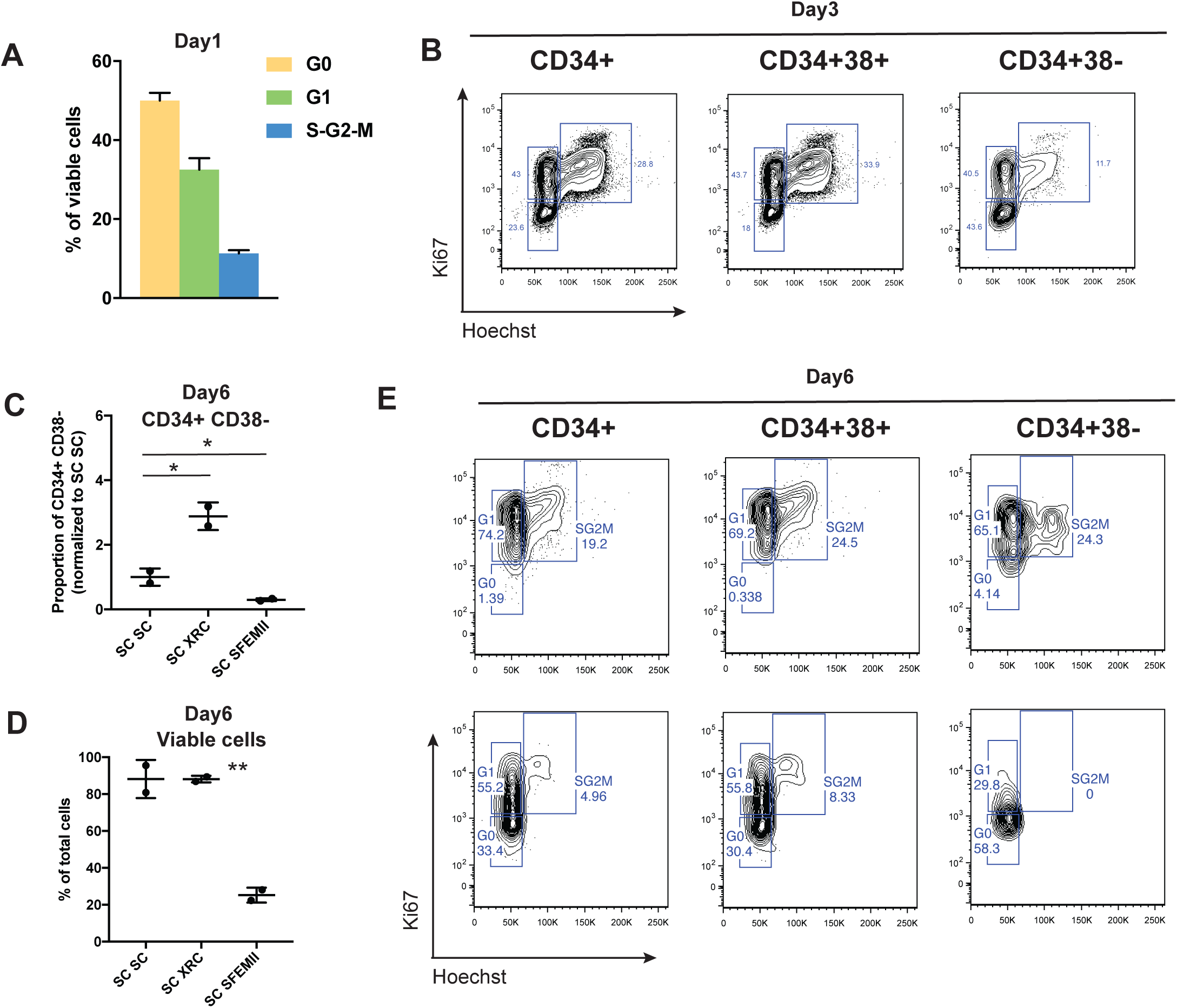
*Related to Figure 4*: Inducing quiescence via XRC treatment after a short period of cycling yields quiescent, primitive HSPC that are distinctive from omission of cytokines. (A) Cell cycle profiles of CD34+ HSPCs at the time of nucleofection (Day1) in Figure 4A. (B) Representative flow plots for cell cycle status 2 days post electroporation of CD34+, CD34+CD38+ (progenitors), and CD34+ CD38- (EE HSPCs). (C) Percentages of EE HSPCs in SC SC, SC XRC, and SC SFEMII normalized to SC SC. Mean ± SD shown. *:p<0.05 by unpaired t-test. (D) Percentages of viable cells in SC SC, SC XRC, and SC SFEMII. Mean ± SD shown. **: p<0.01 by unpaired t-test. (E) Representative flow plots for cell cycle status 5 days post electroporation of CD34+, CD34+CD38+(progenitors), and CD34+ CD38- (EE HSPCs).

### KEY RESOURCES TABLE

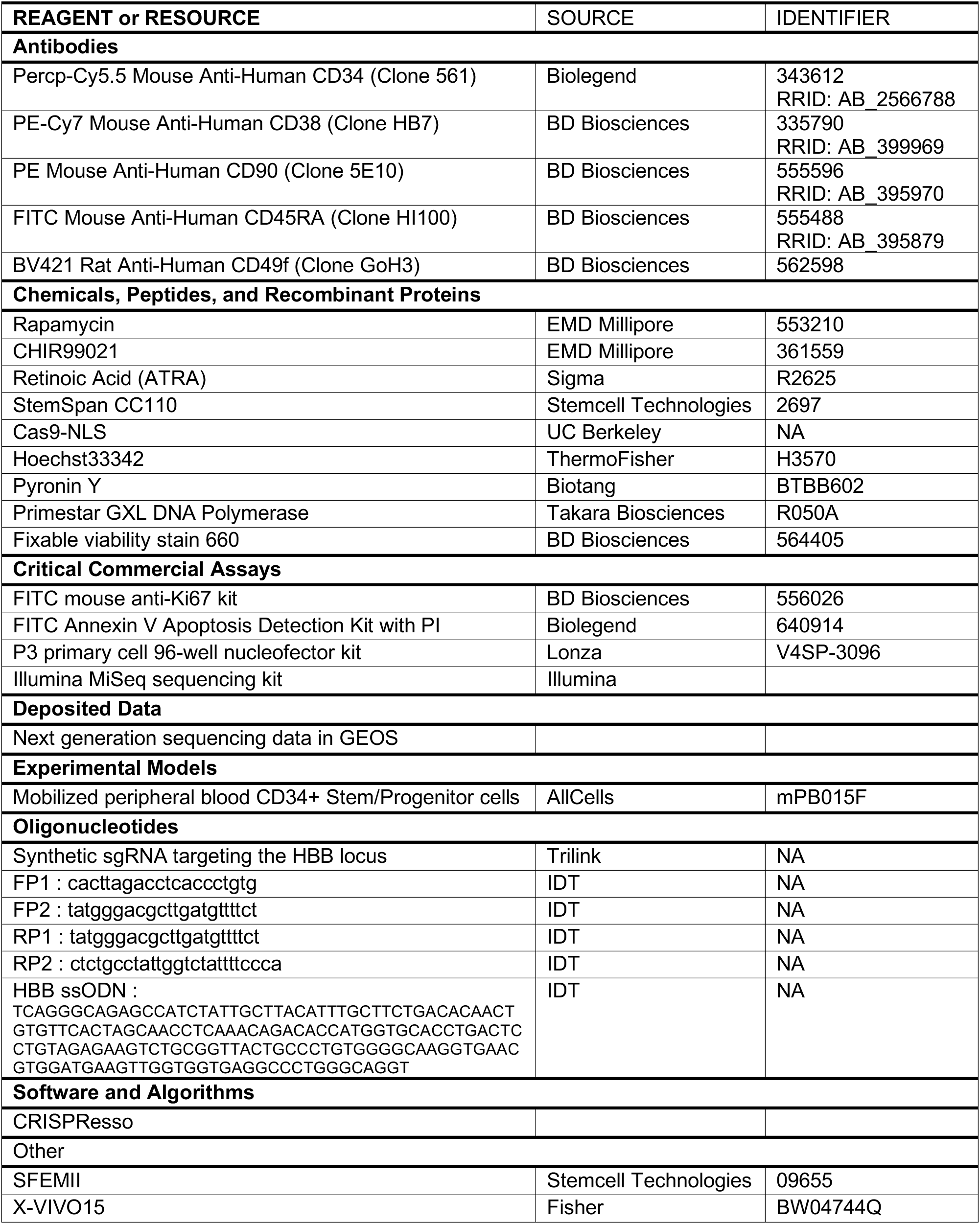

